# Age-dependent decline in sperm quality and function in a naturally short-lived vertebrate

**DOI:** 10.1101/2023.12.13.571459

**Authors:** Silvia Cattelan, Dario Riccardo Valenzano

## Abstract

Maximizing the life-long reproductive output would lead to the prediction that short-lived and fast aging species would undergo no – if any – reproductive senescence. Turquoise killifish (*Nothobranchius furzeri*) are naturally short-lived teleosts, and undergo extensive somatic aging, characterized by molecular, cellular, and organ dysfunction following the onset of sexual maturation. Here, we tested whether naturally short-lived and fast aging turquoise killifish males maximize reproduction and display minimal – if any, reproductive senescence. We analysed age-related changes in sperm traits, the proportion of fertilized eggs, as well as embryo survival. Contrary to the expectation of no reproductive aging, we found that turquoise killifish males undergo extensive reproductive aging, consisting in the age-dependent decline in sperm quality, decreased proportion of fertilized eggs and lower embryo survival. Our results indicate that male turquoise killifish do not trade-off age-dependent soma decline with life-long sustained reproductive fitness. Instead, somatic and reproductive aging appear to occur simultaneously. Our findings question generalized soma vs. reproductive senescence trade-off models and highlight the importance of integrating species-specific ecological and demographic constraints to explain observed life history traits.

## Background

Reproductive senescence consists in the age-dependent decline of reproductive success, which for males depends on the capacity to successfully obtain mates, and to fertilize eggs [1]. Even if a male can mate throughout its life, a decline in the ability to fertilize eggs will negatively affect its reproductive success. Changes in male fertility with age can be investigated by measuring the age-dependent change in sperm traits, and by directly measuring fertilization success and offspring viability [2]. Across animals, evidence of age-related changes in male fertility are mixed, with some taxa showing an overall decline in ejaculate quality as a function of male age, while others showing negligible or even negative reproductive senescence [reviewed in 2].

The variation in male reproductive senescence across species have been suggested to result from trade-offs among life-history traits, i.e. growth, reproduction and survival [3, 4]. Trade-offs emerge from the allocation of limited resources to growth, reproduction and survival, and are predicted to describe most of the variation in life-history strategies along a fast-slow continuum [3, 5]. For instance, species that show continuous growth during life, and thus in which energy is continuously allocated to soma, show negligible or even negative actuarial senescence [6]. A simple but key prediction of soma vs. reproduction trade-off models is that species undergoing extensive somatic aging and high age-related mortality should maximise reproductive output throughout life with negligible – if any - age-related decline in fertility.

Vertebrate species that evolved fast growth, systemic somatic aging, and short lifespan, provide the unique opportunity to test trade-offs between somatic aging and life-long reproduction. The turquoise killifish (*Nothobranchius furzeri*) represents and ideal model to test whether fast growth and somatic aging correlates with high investment in reproduction and no – or negligible – reproductive senescence. Turquoise killifish are among the shortest-lived vertebrates [4-8 months, 7] and have become increasingly adopted in aging research [8, 9]. Turquoise killifish inhabit ephemeral ponds in southeast Africa, where fish mate over a few months [10] by releasing their gametes externally (external fertilization) (**Figure 1**). Upon desiccation of the seasonal ponds, the fertilized eggs manage to survive in the dry soil in a state of developmental diapause [10, 11]. Once the seasonal ponds re-fill with water in the following rainy seasons, the diapausing embryos resume development, and hatch (**Figure 1**). Incubating in dry soil is required for embryos to hatch, leading to a lack of generation overlap from one wet season to the next. Turquoise killifish undergo extensive spontaneous somatic aging, characterized by shorted telomeres [12], cellular senescence [13, 14], brain dysfunction [15, reviewed in 16], cognitive and locomotor decline [17], and eventually a short lifespan [9, 10]. If somatic aging and reproduction trade-off [4, 18], killifish might minimize reproductive aging to overcome their extensive somatic aging.

**Figure 1:**
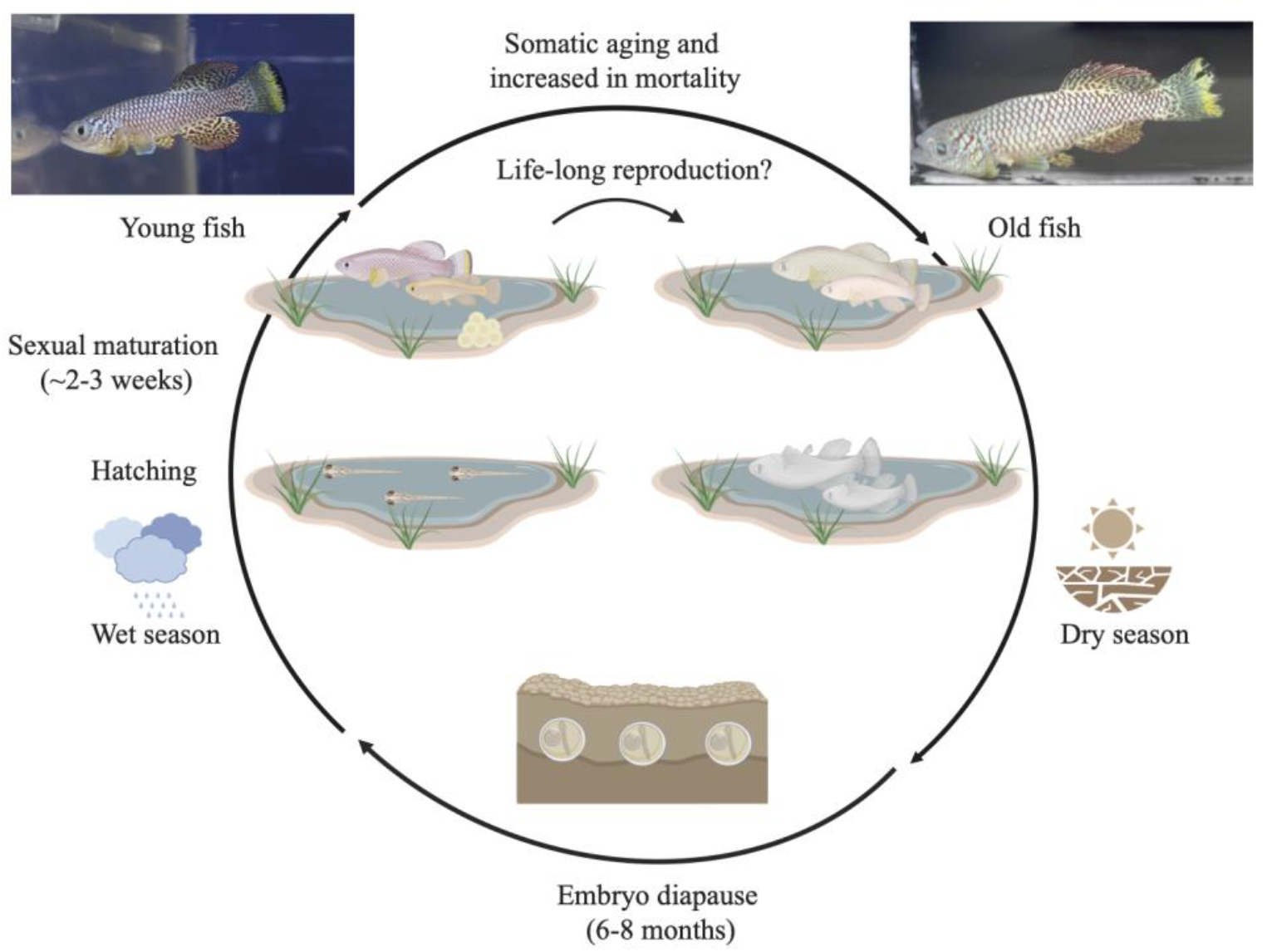
Schematic representation of the life cycle of the turquoise killifish. During the wet season embryos hatch and they quickly reach sexual maturation in 2-3 weeks. After sexual maturation, males and females mate for a few weeks before dying. At the end of the wet season, the ponds dry out and the fertilized eggs remain for months in the dry ground under a state called diapause. On the left an adult (8 weeks) and old (24 weeks) representative GRZ-D males are displayed.

Here, we set out to rigorously test male reproductive senescence in turquoise killifish. Evidence for male reproductive senescence in turquoise killifish males is mixed, with some studies suggesting a deterioration in testes morphology and fertilization capability at very old age [19, 20], while other studies finding limited or no consistent evidence of male reproductive aging [21, 22]. We focus on measuring sperm traits in isolation, and on measuring the impact of male age on egg fertilization and embryo survival. In this work we find that, despite the expectation of limited reproductive aging, male turquoise killifish undergo extensive sperm aging, which impacts fertilization success and embryo survival.

## Results

### Experiment 1: Impact of male age on sperm quality

We investigated the age-related deterioration in the ejaculate by measuring sperm quality parameters that are commonly used in fish [e.g. 23, 24, 25] and in other animals [reviewed in 2]. Specifically, we compared sperm viability, sperm DNA integrity, sperm motility and sperm velocity between adult (9.18 ± 1.87 sd weeks) and old males (22.33 ± 3.99 sd weeks) (**Figure 2a**). After standardizing recent social/sexual history by allowing males to mate for three days, followed by five days of isolation, we dissected the testes from individual males and we released the sperm into Hank’s solution (see **Methods, Experiment 1: effect of age on sperm quality**). Once we isolated sperm from testes, we assessed sperm viability using a live/dead viability Kit for sperm cells in which a membrane-permeant nucleic acid stain (acridine orange) labels live sperm in green, while a membrane-impermeant stain (propidium iodide) labels dead or damaged sperm in red (see **Methods**). We compared sperm viability between 18 adult and 23 old males and found that sperm from adult fish showed a higher proportion of viable cells compared to sperm from old fish (**Table 1, Figure 2b**). On average, the percentage of live sperm over the total was 84.72% (± 1.48 sem) for adult males and 77.3% (± 1.69 sem) for old males.

**Table 1:** Results from the mixed models on the effect of age on sperm traits. All the variables were fitted with a binomial distribution, with the exception of sperm velocity that was fitted with a gaussian distribution.

| Trait | Estimates ± SE | $\chi^2$ | p | Hedges' g (95% CIs) | Number of males |
| --- | --- | --- | --- | --- | --- |
| Sperm viability | $0.48 \pm 0.15$ | 10.991 | $< 0.001$ | 0.99 (0.33, 1.65) | n=18 adult, n=23 old |
| Sperm DNA integrity | $0.31 \pm 0.07$ | 21.454 | $< 0.001$ | 1.77 (0.70, 2.84) | n=9 adult, n=11 old |
| Sperm motility | $0.91 \pm 0.16$ | 33.886 | $< 0.001$ | 1.62 (0.92, 2.33) | n=10 adult, n=13 old |
| Sperm velocity | $-0.77 \pm 2.08$ | 0.136 | 0.712 | 0.22 (-0.44, 0.90) | n=10 adult, n=10 old |

**Figure 2:**
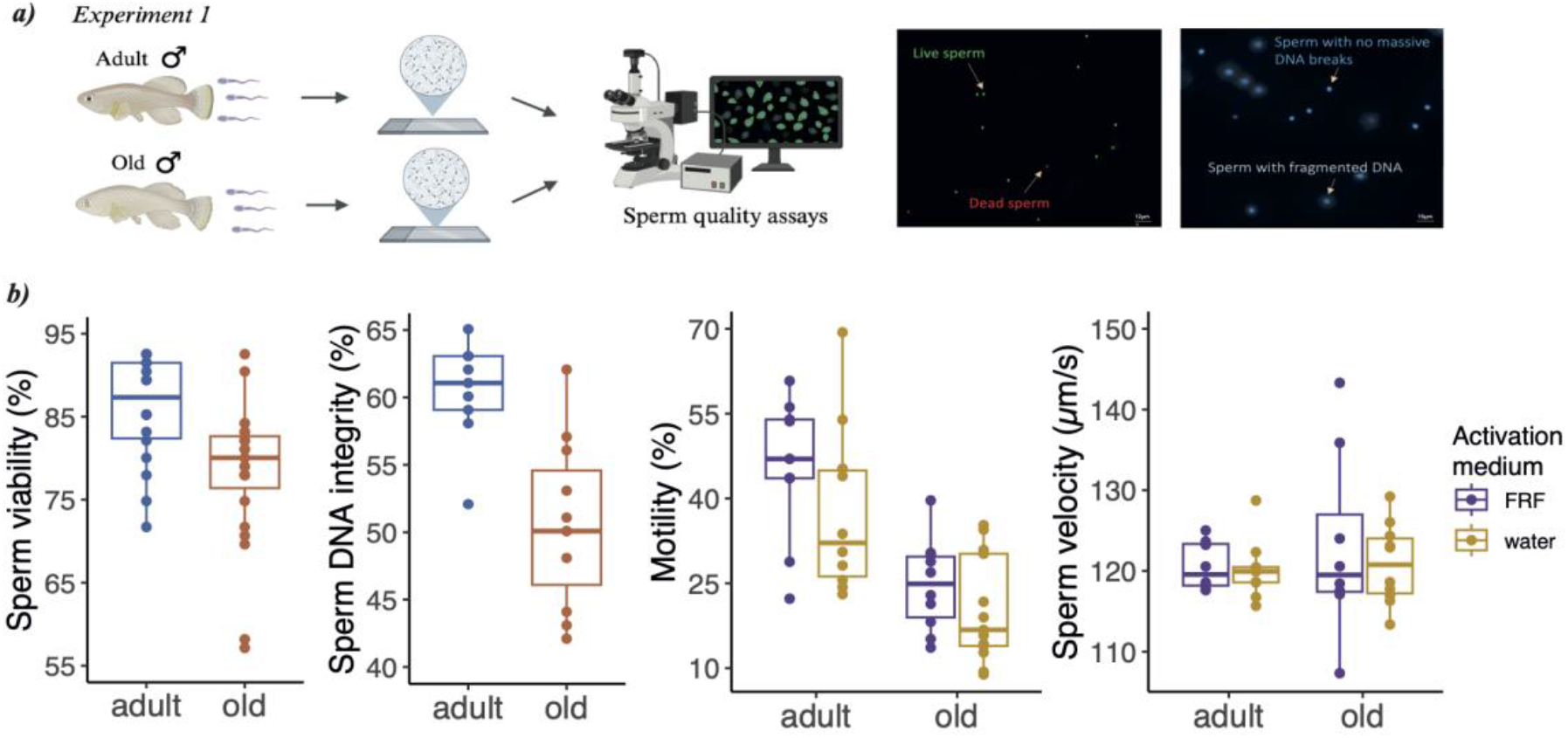
***a)*** Experimental design for Experiment 1 and example of images from the sperm viability (left) and the sperm DNA integrity (right) assays. In the sperm viability assay, live sperm are labelled as green dots while dead sperm as red dots. In the sperm DNA integrity assay, sperm with fragmented DNA appeared as large spots with a blurred halo of chromatin dispersion while sperm with intact DNA appeared as small spots with a compact halo of chromatin dispersion; ***b)*** box plots depicting differences in sperm quality from adult and old turquoise killifish males. All sperm traits, except sperm velocity (p=0.918), are significantly different between adult and old males (p<0.001).

We then set out to test DNA integrity as a measure of sperm quality by employing a sperm chromatin dispersion (SCD) procedure (see **Methods**). We measured DNA integrity in 9 adult and 11 old fish. We found that compared to sperm from old fish, sperm from adult fish had a higher proportion of sperm with intact DNA: 60.33% (± 1.27 sem) vs 50.55% (± 1.88 sem) (**Table 1, Figure 2b**).

We then asked whether sperm motility and sperm velocity, known to be important for fertilization success in many external fertilizer fish species [e.g. zebrafish, 26], differed between adult and old fish. We tested sperm motility and sperm velocity under two conditions: water and female reproductive fluid (FRF), which is the fluid surrounding the eggs of external fertilizers and which in many species can both activate sperm and increase sperm performance [27]. The FRF was collected from 10 female egg batches (age: 8-10 weeks), and the same pool of FRF was used throughout this work (see **Methods**). After sperm activation (either using water or FRF), the sperm solution was transferred on a slide, covered with a coverslip and sperm movements were recorded. Videos captured at 100 frames/second and subsequently analysed using a computer-assisted sperm analyser (CEROS, Hamilton-Thorne).

We compared sperm motility (i.e., proportion of motile cells over the total cells) between 10 adult and 13 old fish and we found that adult males had a significantly higher proportion of motile sperm compared to old males both when sperm were activated with water and with FRF (**Table 1, Figure 2b**). As expected, we found that the activation medium had a significant effect on sperm motility, with a higher proportion of sperm being motile when activated with FRF (34.57% ± 3.4 sem) compared to water (27.89% ± 3.1 sem) (activation medium: **χ**^2^=25.727, *p*<0.001). There was a tendency for sperm from adult males to be more motile in FRF than in water compared to sperm from old males, though this difference did not reach statistical significance (male age * activation medium: **χ**^2^=1.868, *p*=0.17).

While we found that sperm from young individuals were generally more active than from old males, we asked whether sperm velocity (μm/s) – among motile sperm – differed in sperm from adult and old males. Noteworthy, we found no difference in sperm velocity between adult and old sperm, under either activation media (**χ**^2^=0.406, *p*=0.524) (**Table 1, Figure 2b**).

Overall, we found that male age significantly impacts the quality of the ejaculate by decreasing sperm viability, DNA integrity and sperm motility.

### Experiment 2: Effect of male age on fertilization success and embryo survival

To investigate the impact of male age on sperm function, we performed 20 controlled IVFs with sperm from 14 adult (9.1 ± 1.97 sd weeks) and 14 old (26.15 ± 2.54 sd weeks) males and we then measured fertilization success and embryo survival (**Figure 3a**). After standardizing recent social/sexual history and dissecting the testes from individual males as described in Experiment 1, we released the sperm into Hank’s solution. To control for potential differences in egg quality among females, we used eggs from the same females and conducted IVFs using sperm from adult and old males (see **Methods, Experiment 2: effect of age on fertilization success and embryo survival**). We measured fertilization success by counting the fertilized eggs over the total number of eggs used for the IVF (**Figure 3a**). We found a significant effect of male age on fertilization success (**Table 2, Figure 3b**), with adult males fertilizing 81% (± 3 sem) of the eggs and old males fertilizing 65% (± 5 sem) of the eggs.

**Figure 3:**
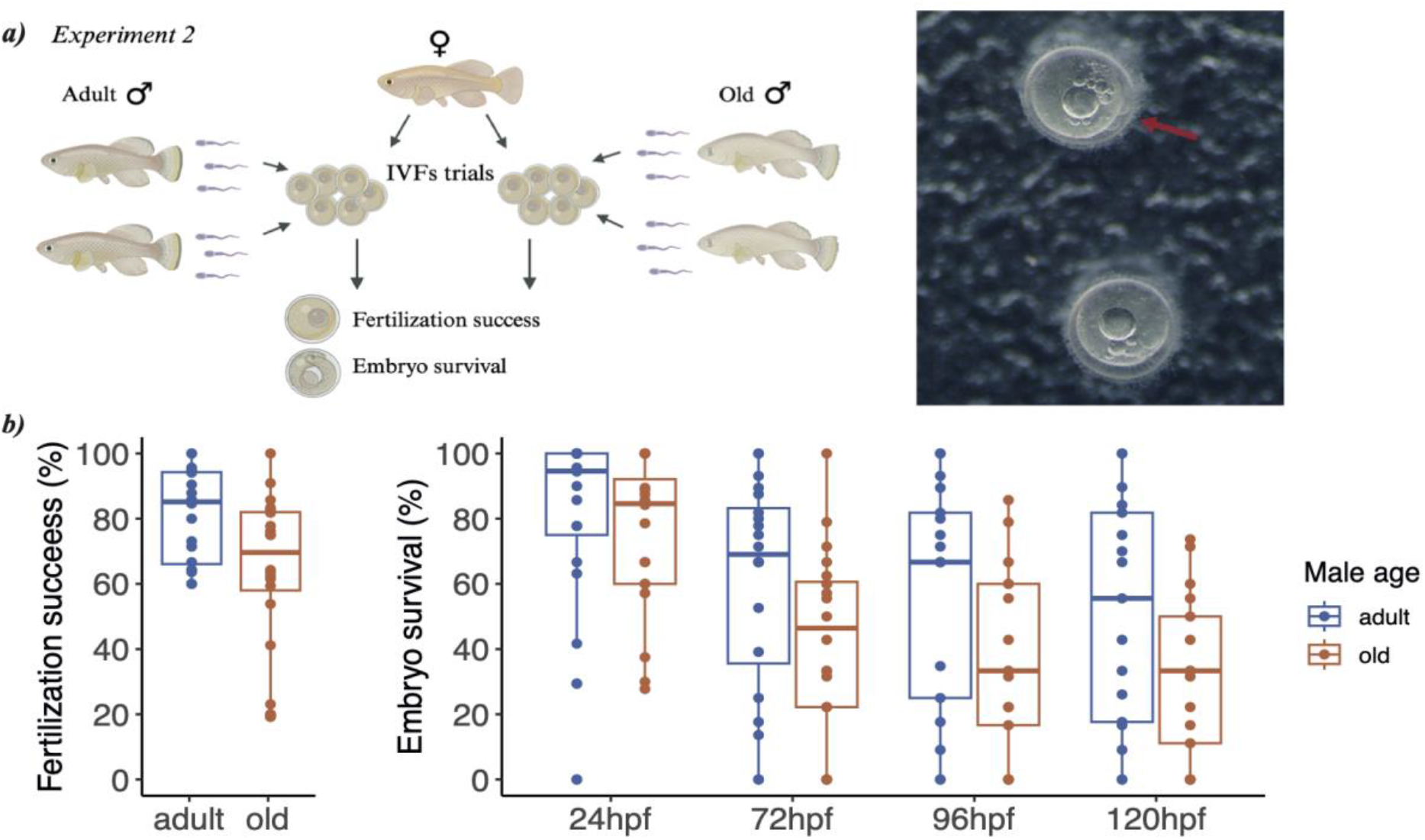
***a)*** Experimental design for Experiment 2 and example of fertilized eggs at 3.5 hours post fertilization. The red arrow indicates the dividing cells; ***b)*** box plots depicting significant differences (p<0.001) between adult and old males in fertilization success and embryo survival over the first five days from fertilization.

**Table 2:** Results from the mixed models on the effect of age on fertilization success and embryo survival. The variables were fitted with a binomial distribution. In total we performed 40 IVF trials, n=20 using sperm from adult males and n=20 using sperm from old males.

| Trait | Estimates ± SE | $\chi^2$ | p | Hedges' g (95% CIs) |
| --- | --- | --- | --- | --- |
| Fertilization success | 1.26 ± 0.20 | 38.556 | < 0.001 | 0.81 (0.17, 1.44) |
| Embryo survival | 0.73 ± 0.13 | 32.062 | < 0.001 | 0.48 (0.15, 0.81) |

We then tested embryo survival by counting the number of live embryos after 24, 72, 96, 120 hours post fertilization (hpf) and calculating the proportion of embryos alive on the total number of eggs fertilized (see **Methods**). We found that embryos fertilized by sperm of old males had a reduced overall survival compared to embryos fertilized by sperm of adult males (**Table 2, Figure 3b**). Embryo survival decreased over time (from 24 to 120 hpf) (**χ**^2^= 166.921, p<0.001), but not differently between sperm from adult and old males (male age*time: **χ**^2^= 2.520, p=0.470).

Overall, we found that male age strongly impacts fertilization rate and embryos survival.

## Discussion

The turquoise killifish is one of the shortest-living vertebrates, with a natural lifespan that ranges between 4 and 8 months, depending on the population [7]. Species showing somatic aging and high actuarial senescence are predicted to maximize the investment in reproduction throughout their lifespan, and thus to show limited or any reproductive senescence. We thus expected to observe a high investment in male fertility throughout the lifespan of turquoise killifish males, by detecting limited sign of male reproductive senescence. Instead, our results point towards the presence of age-related decline in ejaculate quality, which resulted in a strong decline in fertility of old turquoise killifish males. Our results thus question generalized soma vs. reproduction trade-off models, and we suggest alternative processes that could have contributed to the evolution of (male) reproductive aging in turquoise killifish.

Despite the interplay between growth, reproduction, and survival is predicted to explain most of the variation in life-history strategies [4], other processes may explain the evolution of patterns of survival and reproduction [5]. Mutations with no (or slight) effect on fitness in early life but that negatively affect late-life fitness can accumulate in the populations due to the age-related decline of purifying selection, and eventually shorten lifespan [28]. Moreover, demographic history is expected to strongly influence the rate of mutation accumulation. For instance, repeated bottlenecks and habitat fragmentation reduce population size, which in turn decreases the efficacy of selection in removing partially deleterious mutations [29]. In African annual killifishes, limited population size due to repeated bottlenecks and highly fragmented habitats favoured the accumulation of genome-wide deleterious genetic variants [30]. The accumulation of deleterious mutations might have extensively contributed to the evolution of fast somatic aging and short lifespan, both at the species and population level [30, 31]. Hence, demographic constraints and relaxed selection for late-life acting genes, instead of adaptive evolution, appears to have largely contributed to shaping late-life phenotypes in the African annual killifishes. It is thus plausible that the decline in male fertility that we observed does not result from the optimal allocation of resources between reproduction and survival, but rather results from the non-adaptive accumulation of deleterious mutations. Under this scenario, we hypothesise that longer-lived species of annual killifishes, which have accumulated less deleterious genetic variants compared to shorter-lived species [31], should show weaker age-related decline in fertility, compared to shorter-lived species, as the species used in the present study. Future comparative studies on annual killifishes with different lifespan will be instrumental to test further this hypothesis.

While we observed no clear trade-off between reproduction and survival at the species-level, we cannot exclude that trade-offs exist at the population level. Indeed, within the same population, individuals may display alternative life-history strategies, with some individuals reproducing earlier and dying young, and other individuals starting later to reproduce (or having a lower reproductive success) and living longer (the so called pace-of-life syndrome) [32]. For instance, in Soay sheep (*Ovis aries*), normal-horned males have greater annual reproductive success but reduced longevity compared to males with reduced horns [33]. Although, the pace-of-life syndrome has received mixed empirical support, variation in reproductive strategies may be important in shaping life-history variation especially in short-lived species [34]. For instance, *N. furzeri* embryos can either develop fast, by skipping diapause, or can develop slowly by entering diapause and remaining in this state for 5-6 months. Different embryo development strategies may underly differences in life-history strategies during the adult life, with fish skipping diapause having a faster sexual maturation and a shorter lifespan [35]. However, evidence for individual life-history variation linked to diapause is mixed [11] and future studies are required to test for within-population variation in reproductive strategies in the turquoise killifish.

Third, to successfully reproduce, males need to both mate and fertilize the eggs. Traits associated with mating and fertilization success may have independent developmental and aging trajectories. In guppies, for instance, traits associated with mating success (i.e., sexual behaviour and coloration) senesce at a faster rate than traits associated with fertilization success as sperm quality [36], suggesting that different reproductive traits may have different senescence rate depending on the biology of the species. Turquoise killifish males are larger than females, display conspicuous sexual coloration, and intensely court the females; all traits that are supposed to be important to obtain mates. In turquoise killifish traits associated with mating success, such as courtship behaviour and body coloration, may senesce slower than traits associated with fertilization success, thus potentially alleviating the age-related effects of male fertility. However, how traits associated to mating success and fertilization success senesce relative to each other requires further investigation.

Overall, our findings show that turquoise killifish males undergo a spontaneous decline in ejaculate quality, which leads to an impairment of their reproductive functions. Old males fertilized fewer eggs and produce fewer viable embryos. To what extent male age can affect offspring fitness via ejaculate-mediated effects remains an important and open question in evolutionary biology. With age, sperm can accumulate genetic and epigenetic changes [37, 38]. Mutations accumulate in sperm due to the high number of germline cell divisions and can potentially impact offspring health [39]. Similarly, the sperm methylome undergoes age-related changes, which have been linked to the reduced lifespan observed in offspring from old fathers compared to young fathers [40]. To assess the evolutionary consequences of male reproductive aging, future work is needed to investigate male age effects on offspring fitness. Species undergoing spontaneous reproductive senescence, such as the turquoise killifish, will help clarify the impact of male reproductive aging on offspring fitness.

## Methods

### Fish maintenance

The turquoise killifish used in this study were laboratory-reared fish from GRZ-D strain maintained at the Fish Facility of the Leibniz Institute on Aging (Jena, Germany) under standard laboratory conditions (12:12 light–dark cycle; water temperature 26 ± 1 °C; pH 6.5-7.5). Full details on housing and breeding conditions are reported in [41]. The median lifespan of the fish cohort used in the experiments was 23 weeks (22.82-24.00, 95% CIs).

### Experiment 1: effect of age on sperm quality

Before sperm traits analysis, each adult male was allowed to interact and mate with at least 2 females in a 3.2L tank (Acqua Schwarz) to standardize recent social/sexual history. To allow the replenishment of sperm reserves, after 3 days of mating, each male was isolated from females for 5 days [42]. At day 8, each male was euthanized, testes were removed and placed into a 500μL of Hank’s buffer solution and maintained in ice [43]. To allow the release of sperm into Hank’s solution, we spun back and forth the testes in the tube for 1min using forceps, following an established protocol [44]. Immediately thereafter, the testes were removed from the tube. All sperm assays (sperm motility and velocity, sperm viability, sperm DNA integrity) were performed within 1h from sperm collection (**Figure 2a**). The average age of adult males was 9.18 ± 1.87 weeks (mean ± sd) and that of old males was 22.33 ± 3.99 weeks, which corresponds to the median lifespan in this cohort population. Details on the number of adult and old males used in each sperm assay are reported in **Table 1**.

### Sperm motility and velocity

For each sperm motility and velocity assay, sperm were activated using either distilled water or the female reproductive fluid (hereafter FRF). FRF surrounds the eggs of external fertilizers and in many species it can both activate the sperm and increase sperm performance [27]. FRF was collected by 10 females of 8-10 weeks. Females were squeezed from the eggs following an established protocol [42] and FRF was collected using a 3μL pipette (Drummond) and transferred to a tube, which was then maintained at -80C until the experiment was run. On the day of the experiment, FRF from different females was pooled and then diluted to final concentration of 5% in distilled water. One μL of sperm solution was activated with either 2.5μL of distilled water or with 2.5μL of diluted FRF on a 12-well slide (MP Biomedicals) coated with a 1% polyvinyl alcohol to prevent sperm from sticking to the glass slide [25]. The slide was immediately covered with a coverslip (22×22 mm) and sperm motility was recorded through a digital camera connected to a phase-contrast microscope (Axiovert 25 clf inverted microscope, Zeiss). Videos were captured at 100 frames/second and subsequently analysed using a computer-assisted sperm analyser (CEROS, Hamilton-Thorne). We measured sperm motility (i.e., proportion of motile cells over the total cells) and sperm velocity (VCL, μm/s). These parameters are known to be important for fertilization success in many external fertilizer fish species [e.g. zebrafish, 26]. We recorded sperm motility parameters on an average of 102.48 ± 15.16 (mean ± SE) sperm cells per male.

### Sperm viability

Sperm viability was measured using a live/dead viability Kit for sperm cells (Halotech DNA) [e.g. 25]. A membrane-permeant nucleic acid stain (acridine orange) labels live sperm in green, while a membrane-impermeant stain (propidium iodide) labels dead or damaged sperm in red. Six μL of each sperm solution were placed on a microscopic slide and 0.8μL of acridine orange and 0.8μL of propidium iodide were gently added. After covering the final solution with a coverslip (24×30 mm), fluorescent images of samples were taken at 40X magnification using a fluorescent microscope (Axiovert 25 clf inverted microscope, Zeiss). From digital images, we counted the number of live (green) and dead (red) spermatozoa (see **Figure 2a**). For each male, we analysed an average of 261.10 ± 14.95 (mean ± sem) total sperm cells.

### Sperm DNA integrity

We assessed the proportion of sperm with intact DNA using the Halomax-SCD kit (Halotech DNA), which is based on the sperm chromatin dispersion (SCD) procedure. We diluted sperm sample in Hank’s buffer solution at the recommended concentration and we then followed the manufacturer’s instructions. To visualize the sperm DNA, each well on the slide (8 wells/slide) were stained with DAPI staining (1:1000 dilution) and placed under a fluorescent microscope (ApoTome, Zeiss). Sperm presenting fragmented DNA appeared as large spots with a blurred halo of chromatin dispersion while sperm with intact DNA appeared as small spots with a compact halo of chromatin dispersion (see **Figure 2a**). We counted an average of 273.65 ± 12.76 (mean ± sem) total sperm cells per male.

### Experiment 2: effect of age on fertilization success and embryo survival

We investigated the impact of male age on fertilization success and embryo survival by performing controlled IVFs in which sperm from adult and old males were used separately to fertilize the eggs from the same female (**Figure 3a**). For this experiment, we used 28 males (n=14 adult, n=14 old). The average age of adult males was 9.1 ± 1.97 weeks (mean ± sd) and that of old males was 26.15 ± 2.54 weeks. To standardize socio/sexual conditions and to collect sperm we followed the same protocol described in Experiment 1. After testes collection (see **Methods, Experiment 1: effect of age on sperm quality**), we counted sperm number using a cell counter (TC20, BioRad). To control for differences in sperm number among males, we standardized sperm concentration of males used in the same IVF trial. We then pooled the sperm from two males of the same age to reduce the potential for specific male x female effects. To perform IVFs, we used in total 20 females. Eggs were collected from 20 females (age range: 13-21wph) following an established protocol [42] and incubated in a bovine serum albumin (BSA) solution at 4% [42] in order to maintain the eggs in the inactivated state. Sperm and eggs were used fresh, i.e. within 30 minutes, after collection. The eggs of each female were split in two batches and placed in two different petri dishes using a disposable Pasteur pipette. To control for potential differences in egg quality among females (e.g., due to age or intrinsic quality), we used each batch of eggs from the same female to perform an IVF with either the sperm pool from adult or old males, for a total of 40 IVFs (**Figure 3a**). Before IVF, we removed the BSA from the eggs using a disposable Pasteur pipette, we activated pooled sperm with 200μl of water and we released the sperm solution over the eggs. After 3 minutes from fertilization, we incubated the eggs in a methylene blue solution (1:1000) and incubated the eggs at 28°C. To assess fertilization success, we checked eggs after 3.5 hours (**Figure 3a**), after which we removed the unfertilized eggs. To score embryo survival, we checked fertilized eggs after 24, 72, 96, 120 hours post fertilization (hpf).

### Statistical analyses

We conducted data analyses using R v 4.2.0 [45]. We fitted mixed effect model using ‘lme4’ package [46]. We tested sperm viability, sperm DNA integrity, sperm motility, fertilization success and embryo survival by fitting a generalized mixed model (‘glmer’ function) assuming a binomial error distribution and a logit link function. We instead tested sperm velocity by fitting a linear model with the ‘lmer’ function. We fitted all models with male age (adult vs old) as a fixed effect. In the models for sperm motility and sperm velocity we also fitted the activation medium (level: water, FRF), and in the model for embryo survival we fitted the time after fertilization’ (level: 24hpf, 72hpf, 96hpf, 120hpf) as a fixed factor. To control for potential batch effect in sperm traits, we fitted models using the male block as random factor. The models on fertilization success and embryo survival were fitted with i) female identity, as the eggs from the same female were used in two IVF trials (either with sperm pool from adult or old males), and with ii) sperm pool identity, to account for the fact that the same sperm pool were used in different IVF trials. For each linear model, we checked the assumption of normality of the residuals by inspecting residuals vs. fitted values. For each generalized mixed model, we tested for overdispersion by using the ‘dispersion_glmer’ function within the ‘lme4’ package. The model on sperm viability was overdispersed (2.10) and we corrected it by adding an Observation-level random effect. We calculated *p* values of the fixed effect by Type II Wald chi-square tests using the ‘Anova’ function in the ‘car’ package [47]. The effect sizes (Hedges’ *g*) and the associated 95% CIs were calculated using the ‘effectsize’ package [48].

## Ethics statement

The research was performed under the EU laws (EU Directive 2010/63/EU). All applications for the use of animals were approved by the regional authorities for animal welfare (Thüringer Landesamt für Verbraucherschutz-TLV) based in Germany. Fish were euthanized under the approved license O_DV_21-24 by immersion in a 1.5 g/L Tricaine solution.

## Author contributions

S.C. and D.R.V. conceived the study, S.C. performed the experiments and analyzed the data, S.C. and D.R.V. wrote the manuscript.

## Acknowledgements

We thank the Fish Facility staff at the Leibniz Institute on Aging, especially Beate Hoppe, Uta Naumann, Simone Dunkel, Christin Hahn, Johannes Wilfert, Clemens Peters, Ronja Baal, and Marcel Münnich, and the Imaging Facility staff for providing constant support to our work. We thank Kathrin Reichwald for providing support with ethical permissions. Finally, we thank the Evolutionary Biology research unit at the University of Padova for the use of the computer-assisted sperm analyser (CASA) and especially Alessandro Devigili for providing technical support with the CASA. This work was supported by the Leibniz Institute on Aging.

## Competing interests

The authors declare no competing interests.

## Notes

### Competing Interest Statement

The authors have declared no competing interest.

### Summary of Updates

The introduction was revised to improve clarity. The result section was updated by including new results on the effect of FRF fluid on sperm motility. A figure regarding the life cycle of the turquoise killifish was included (new Figure 1).

